# Extension of a *de novo* TIM barrel with a rationally designed secondary structure element

**DOI:** 10.1101/2020.10.16.342774

**Authors:** Jonas Gregor Wiese, Sooruban Shanmugaratnam, Birte Höcker

## Abstract

The ability to construct novel enzymes is a major aim in *de novo* protein design. A popular enzyme fold for design attempts is the TIM barrel. This fold is a common topology for enzymes and can harbor many diverse reactions. The recently published *de novo* design of a four-fold symmetric TIM barrel provides a well understood minimal scaffold for potential enzyme designs. Here we explore opportunities to extend and diversify this scaffold by adding a short *de novo* helix on top of the barrel. Due to the size of the protein we developed a design pipeline based on computational *ab initio* folding that solves a less complex sub-problem focused around the helix and its vicinity and adapt it to the entire protein. We provide biochemical characterization and a high-resolution X-ray structure for one variant and compare it to our design model. The successful extension of this robust TIM-barrel scaffold opens opportunities to diversify it towards more pocket like arrangements and as such can be considered a building block for future design of binding or catalytic sites.

## Introduction

The tight coupling of protein structure and function motivates the field of protein design to pursue the construction of modified or novel protein functions such as enzyme catalysis, signaling, binding and many more. The design of enzymes is of particular interest due to their applicability but also because it poses a thorough test for our understanding of what defines activity and selectivity. An intensively studied protein fold is the TIM or (βα)_8_-barrel. Enzymes with this fold are ubiquitously found among organisms and efficiently catalyze a wide variety of reactions (Wierenga et al., 2001; Sterner and Höcker, 2005). The fold is composed of an eightfold repeat of βα-units, with eight parallel β-strands forming a circular sheet in the core that is surrounded by the eight α-helices. While the “bottom side” of the barrel with its αβ-loops provides stability to the barrel, the “upper” part including the βα-loops usually contains the catalytic function of TIM-barrel enzymes. The twisting central β-sheet provides a cavity at the opening of the barrel that is often used as substrate binding site in combination with extensions at the βα-loops. These extensions also often play an important role in positioning catalytic residues and shielding the catalytic site from solvent. Due to these exceptional characteristics of TIM barrels they are highly attractive targets for enzyme design approaches. In fact, naturally occurring TIM barrel have already been modified to turn over non-natural substrates, e.g. carrying out a retro-aldol reaction (Jiang et al., 2008). Computational approaches for the design of novel enzymes have been progressing in recent years and their combination with directed evolution has already proven very successful for some reactions (Lechner et al., 2018).

However, the ability to design an enzyme from scratch including its tailor-made scaffold is still a major aim that might be tackled with computational methods for structure prediction. Tools exist for structure prediction that can be used for its inverse, the protein design problem, to predict the structure of amino acid sequences. The widely used Rosetta molecular modeling suite provides tools for both problems. Its *ab initio* structure prediction algorithm generates fragment libraries for a target sequence from the protein structure database and searches conformational space using a Monte Carlo procedure (Bonneau et al., 2001; Simons et al., 1997). The protein design algorithm also uses Monte Carlo optimization to populate a given backbone with energetically optimal residues and uses structure evaluating energy functions for scoring (Kuhlman and Baker, 2000). While the suite proved its utility in several CASP competitions (Bonneau et al., 2001; Raman et al., 2009), it was also the core design tool for the first *de novo* TIM barrel with four-fold symmetry (Huang et al., 2016). This design aimed at providing an idealized TIM barrel with minimal loops, formed from a four-fold repeated sequence. The design called sTIM11 is a monomeric and stable protein that with its X-ray structure provides a suitable starting point for further enzyme design approaches.

In this work we made the first step towards that goal by extending sTIM11 by a simple structural element, namely an additional helix that we inserted in a loop at the upper part of the barrel. We show with a high-resolution X-ray structure that the core protein is properly folded and the extension is formed while the protein’s biophysical properties are maintained. This *de novo* helical extension can be considered as a structural building block for further design approaches towards binding and catalytic sites by providing additional structural mass at the minimal sTIM11 barrel surface.

## Results

### Scaffold and target site identification

The minimal sTIM11 barrel provides a fairly flat top surface with seven adjoining βα-loops pointing into the region that typically comprises the active site of TIM-barrel enzymes (Huang et al., 2016). These loops were considered most suitable to introduce additional functionalities and extensions. Since the introduction of a secondary structural element might interfere with the folding process we considered carefully where to insert the extension. Since the two halves of sTIM11 can assemble into a TIM-barrel like dimer, we believed the loop in the middle of the protein sequence to be maximal tolerant for the structural modification and chose it as the target site for the helical extension (**Figure 1**). We further introduced a small change to the sTIM11 scaffold by removing two cysteine residues that were originally introduced to form a disulfide bridge to improve barrel closure (Huang et al., 2016). However, this disulfide bond did not form, so we removed the two cysteines to generate a cysteine-free variant named sTIM11noCys whose characteristics and crystal structure are reported in Romero-Romero *et al*. (2020).

**Figure 1:**
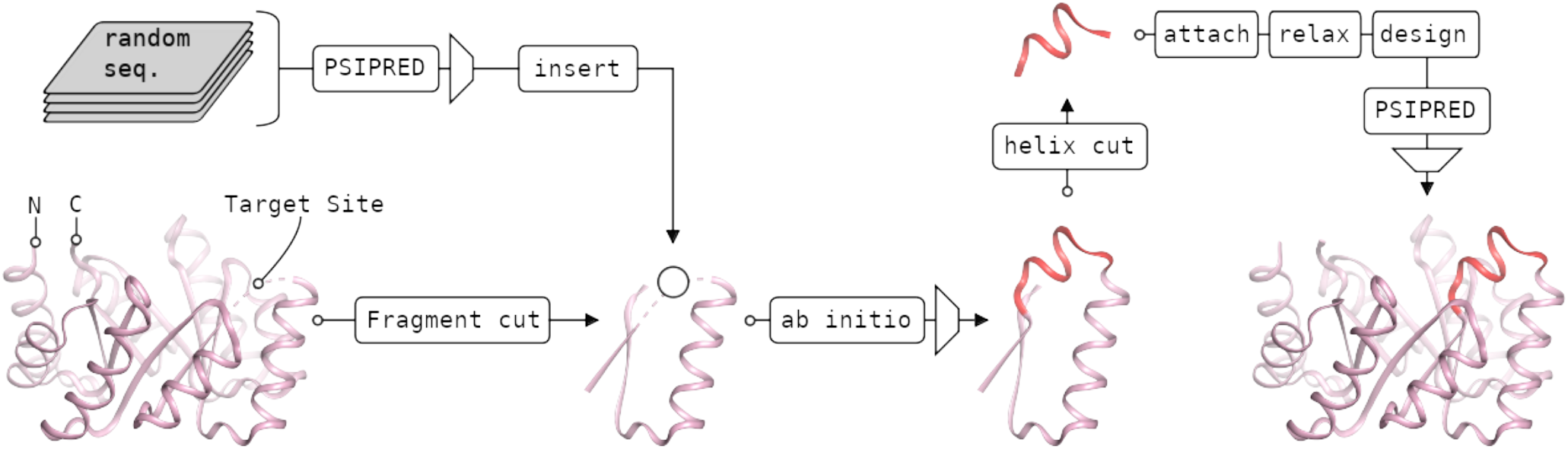
Design pipeline. A βαβ-fragment was extracted from sTIM11, whose structure could be predicted reliably and independently from its context. The target loop (circle) was substituted by random sequences of variable lengths that were predicted to fold into the desired helical structure by PSIPRED. Then the structures were predicted using Rosetta and highly scored models were used further. The α-helix extension (red) was transferred back to the full barrel and adapted to the changed structural context by a Rosetta design step. Only those candidates were used for experimental investigation, that were still predicted to adapt the correct secondary structure according to another PSIPRED run.

### Design pipeline

Our aim was to extend sTIM11noCys by an arbitrary α-helix without any predefined backbone and thus the following design pipeline emerged (**Figure 1**). We wanted to design the model by structure prediction using the corresponding Rosetta *ab initio* method. However, the protein comprises 184 amino acids and thereby surpasses the sequence length limit for reliable folding results. Therefore, we simplified the problem by reducing the folding target to a smaller fragment of the barrel including the surroundings of the target site for insertion. A βαβ-fragment of only 31 residues was found to fold into its tertiary structure independently from the full barrel context when using the Rosetta structure prediction method. In order to find insertions that were likely to fold into an α-helix, more than 150,000 random amino acid sequences of 7 – 14 residues were generated and their secondary structure was predicted using PSIPRED (Jones, 1999). The predictions were filtered for α-helical content and sorted by their prediction score to select the top 30 candidates. These sequences were inserted into the target barrel fragment, thus obtaining the final sequences for the actual Rosetta *ab initio* structure prediction experiments. 1000 models were generated per candidate and multiple sequences were found to fold *in silico* into the desired tertiary structure, thereby recovering the barrel fragment with an additional α-helix at the target site. The structures were sorted by RMSD when superimposing the βαβ-fragment onto the parental barrel and for the top 12 candidates, the α-helix was manually transferred to the full barrel. An additional Rosetta relax step was performed to smooth introduced structural deviations from the manual intervention.

Since the promising tertiary structure had been found in the context of the fragment and not in the context of the entire barrel, the residues of the helix had to be optimized locally to fit with adjacent residues. Therefore, about 50 % of the residues of the α-helix extension was allowed to mutate freely in a subsequent Rosetta design step. Once again, those sequences were filtered by another PSIPRED step to raise the chance of obtaining the desired secondary structure. Four candidates were chosen manually based on hydrophobic packing and predicted polar contacts for experimental characterization. While all candidates expressed and indicated properly folded proteins (**Supplemental Figure 1**), only one candidate, sTIM11_helix3, yielded diffracting crystals for which an X-ray structure could be solved.

### Biochemical evaluation of sTIM11_helix3

After cloning the designed sequence, the corresponding protein sTIM11_helix3 was expressed and purified. Subsequent analytical size exclusion chromatography of the purified protein revealed a single peak, corresponding to a homogeneous monomeric species (**Figure 2**). Circular dichroism spectra were compatible with folded protein containing a mixed α-helical and β-sheet content comparable to sTIM11noCys. Moreover, the design showed a similar thermal stability to sTIM11noCys with a melting temperature of 66 °C.

**Figure 2:**
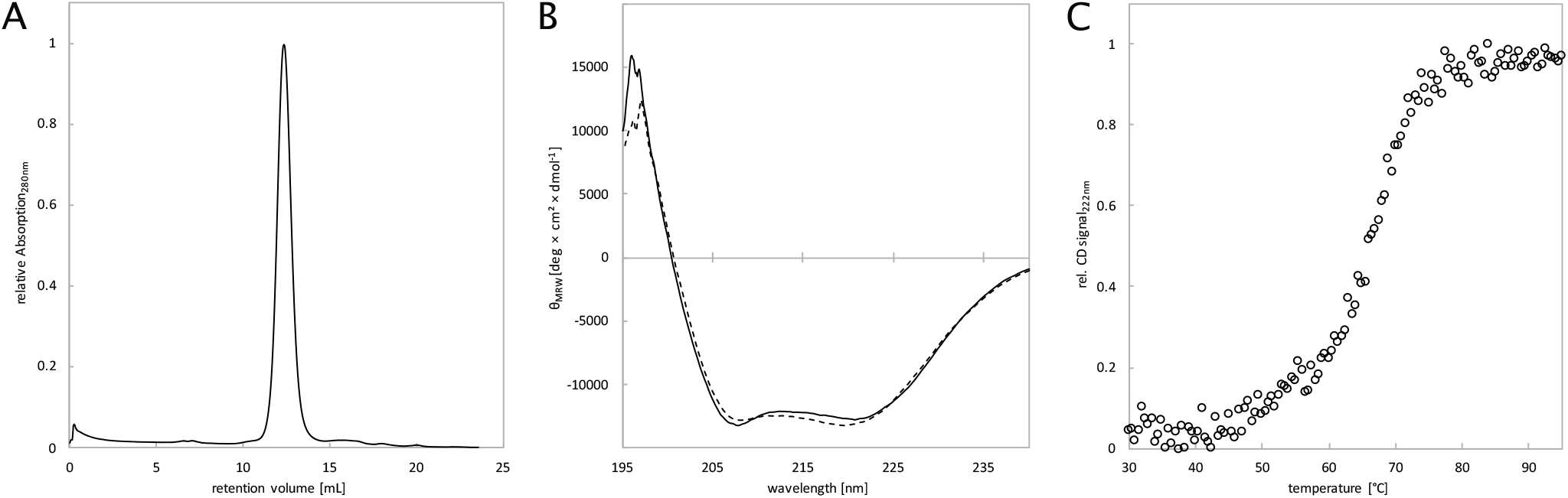
Biochemical Characterization of sTIM11_helix3. (A) Analytical size-exclusion chromatography, (B) circular dichroism measurements before heating (solid line) and after cooling down (dashed line) and (C) thermal melting upon heating.

### Structural evaluation by X-ray crystallography

The protein was crystallized and diffraction data collected and processed to a resolution of 1.58 Å (**Table 1**). The X-ray structure was solved by molecular replacement with four copies of a quarter barrel of the four-fold symmetric sTIM11 (PDB: 5BVL). The B-factor distribution of the final structure revealed highly reliable areas for the core of the TIM barrel, which is clearly recovered in the structure. The introduced helix showed elevated B-factors, making it more difficult to model. Due to the increased flexibility in the region of interest we refined our structure using a feature enhanced map (FEM), reducing the level of noise and model bias (Afonine et al., 2015). The obtained map allowed us to reliably build the entire structure, making the introduced helix clearly observable so it can be compared to the theoretical design model.

**Table 1.**
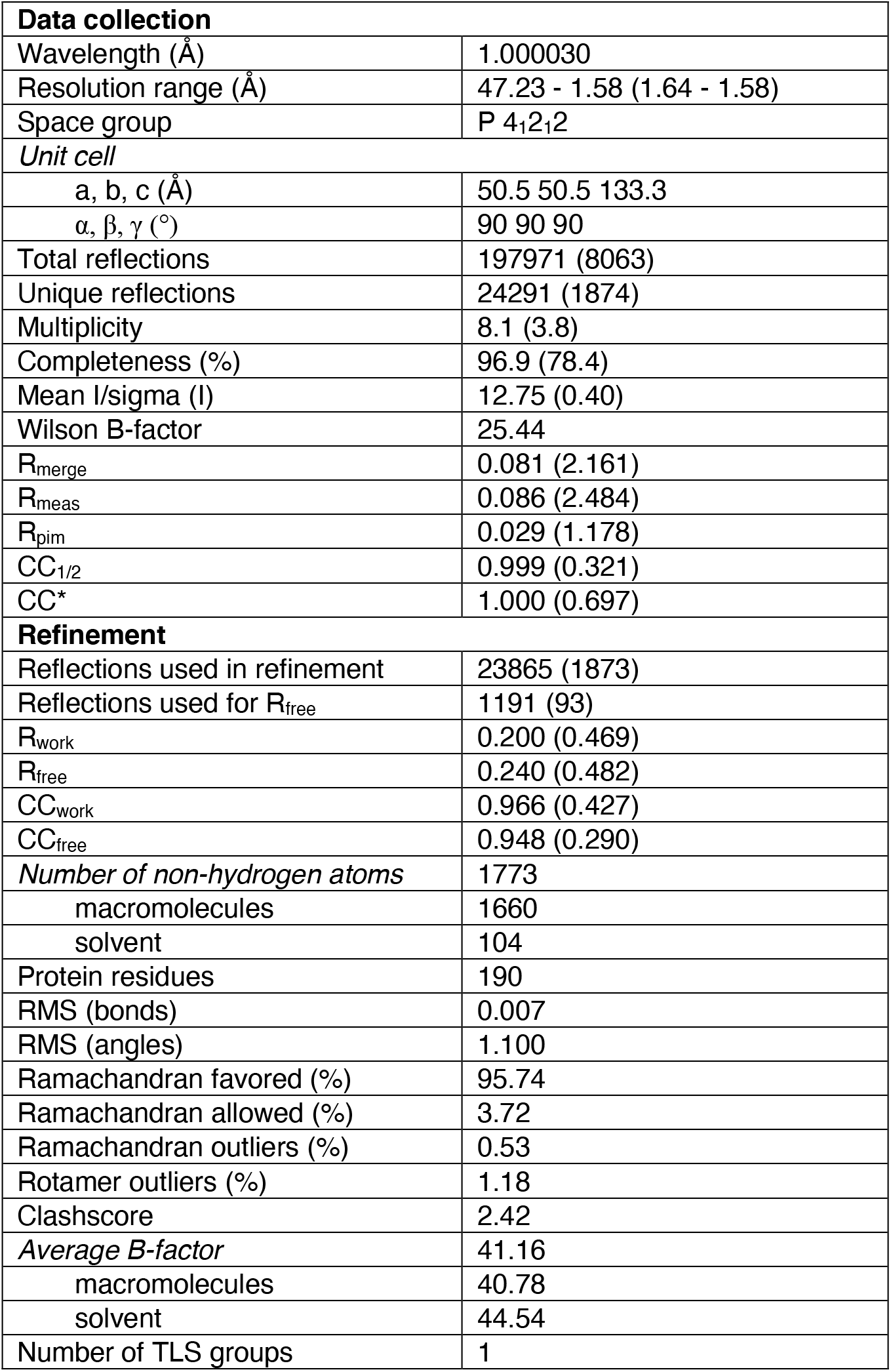
Data collection and refinement statistics. Statistics for the highest-resolution shell are shown in parentheses.

### Comparison of design model and X-ray structure

Superposition of the design model and the X-ray structure show that most parts of the structures are identical. In the core TIM barrel, only the polypeptide chain subsequent to the target site of extension showed structural deviations (**Figure 3**). But also, the helical extension showed a small but significant deviation. While a helical structure was formed, it is wound slightly different than expected and forms a 3^10^ helix instead of a classical α-helix. In addition its position is slightly twisted and it packs more closely and flat onto the barrel surface compared to the design model, which projected the helix to be more exposed. In fact, the design model showed Met94 and Ala95 to dock into a space at the barrel surface without any major structural changes to the main barrel. Interestingly, the experimental structure revealed a different hydrophobic packing and revealed Met94 to rather enter the barrel by inserting its side chain in between the β-sheet core and the outer barrel α-helix (**Figure 4**). Due to this, several polar contacts that had been predicted between the first residues of the inserted helix and its surrounding could not form, but a polar contact between Ser90 and Asp71 is established instead. The following loop that connects the extension with the subsequent α-helix is initiated by Pro98 as designed. But the length of the loop turns out to be longer and more flexible than expected due to a slight unravelling of the barrel helix. In the X-ray structure the loop is solvent exposed and numerous water molecules make polar contacts with its residues. The slightly shortened TIM-barrel helix subsequent to the extension also deviates from its usual position and orientation. It is shifted downwards. Thus, the *de novo* TIM barrel accommodates the newly inserted 3^10^ helix by a rearrangement of the packing of the barrel helix. The snug interaction of the inserted short helix provides a likely explanation for the observed structural deviations of the TIM-barrel core helix.

**Figure 3:**
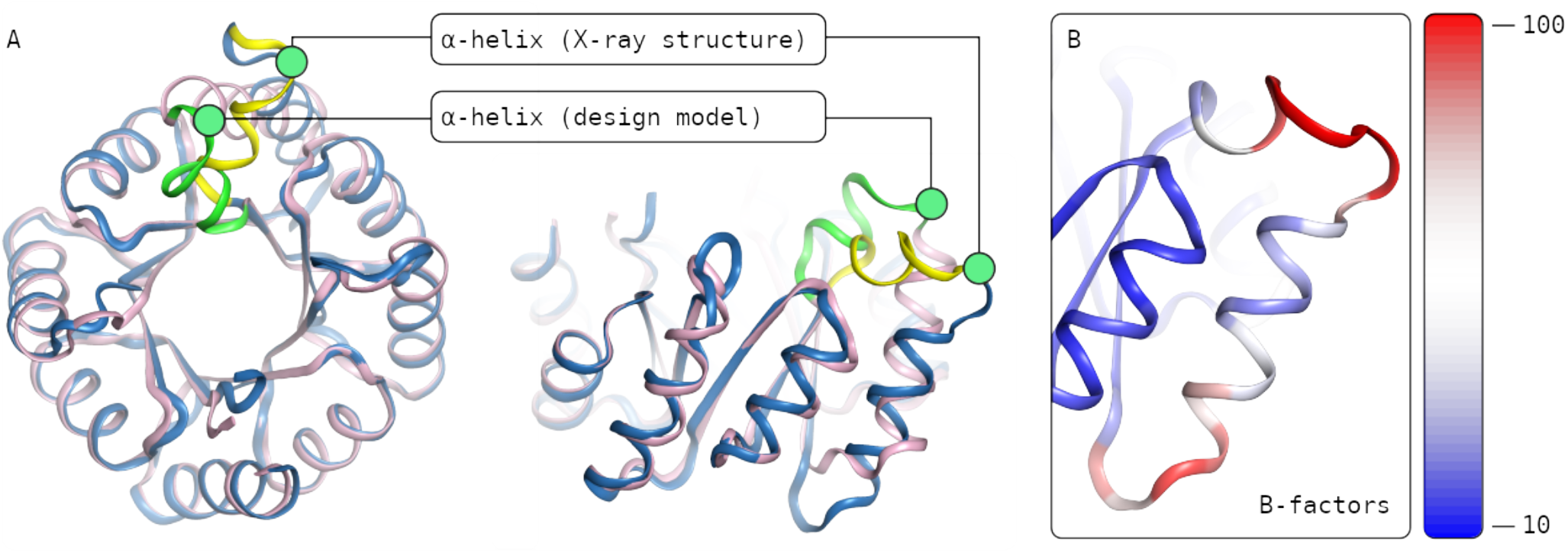
Characterization of the X-ray structure in comparison with the design model. The superposition of the X-ray structure (blue) with the design model (purple) in a top down view (A, left) reveals a close to identical fit. The region of extension shows some structural difference, the inserted α-helix in the X-ray structure (yellow) is twisted in comparison to the prediction (green). The side view (A, right) illustrates that the α-helix is in a different position than predicted and that the following helices within the barrel spatially deviate as well. The B-factor coloring (B) points at the apparent high flexibility in the region of interest.

**Figure 4:**
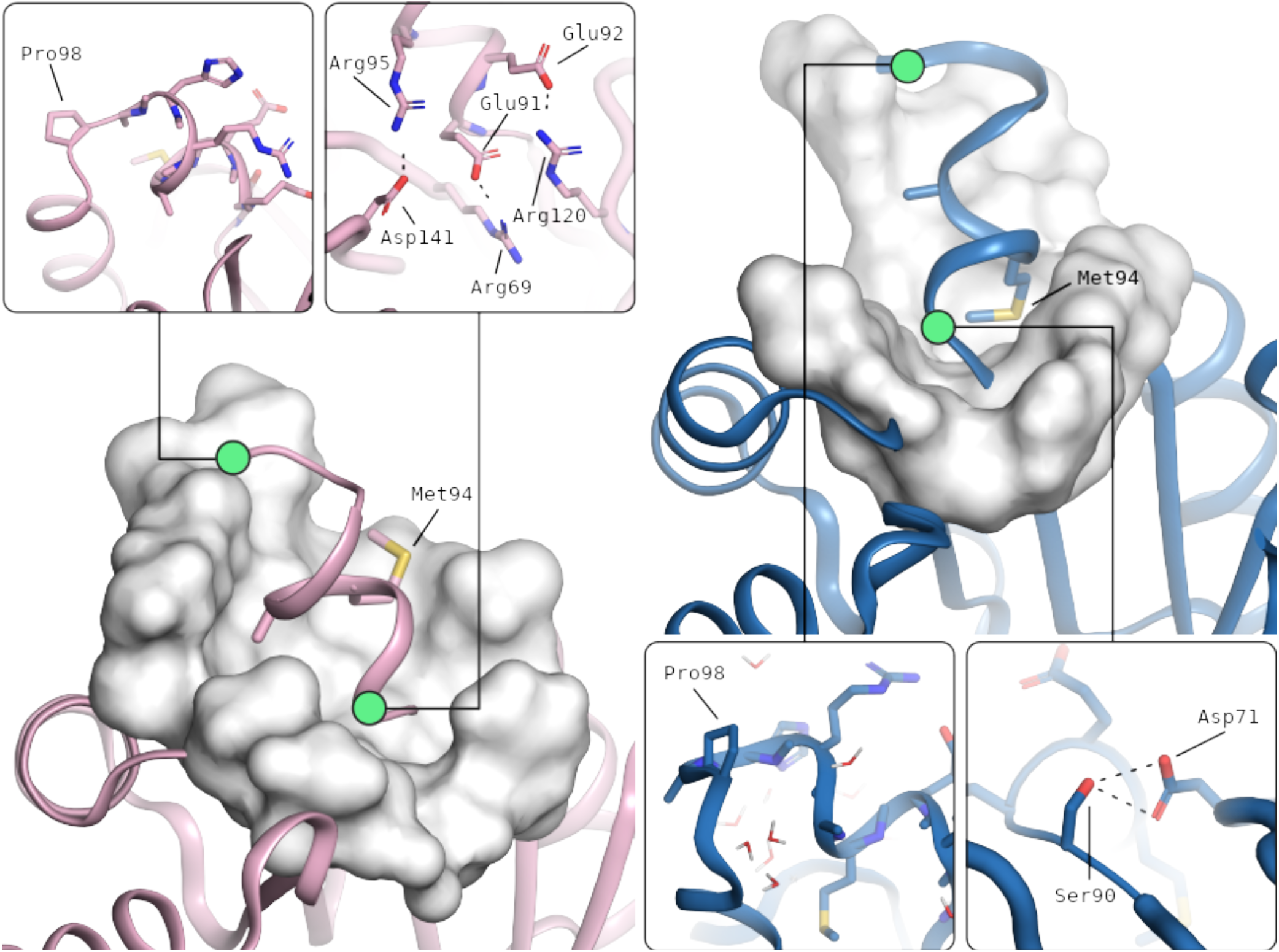
Features of the α-helix in the design model (left) and in the X-ray structure (right). Hydrophobic residue interactions differ (large scale images). In the design hydrophobic residues are packing flat onto the top of the barrel, while in the X-ray structure Met94 rather pierces the top of the barrel in between β-sheet and α-helices. The helix breaking Pro98 (small scale images, left) resulted in an enlarged loop compared to the design which is surrounded by water in the X-ray structure and polar contacts (small scale images, right) are less prominent and differently formed in the X-ray structure.

## Discussion

In this work we provide a first structural extension to the *de novo* TIM barrel sTIM11. Instead of attaching a natural occurring fragment to the barrel, we decided to introduce a short additional α-helix connected by short loops. This required careful integration to the residue environment of the barrel by comprising various computational tools for sequence screening and optimization. By not constraining the design target at the beginning to any fixed backbone, we could not use residue optimization methods initially but performed the first design step by using structure prediction methods instead. We found it to be sufficient to work with just a structural fragment of the barrel in order to find the desired structural extension in a randomly sampled sequence space. After adapting the variants to the adjacent residues in their barrel environment using Rosetta design, well-folded proteins were obtained and the successful extension of one design could be confirmed experimentally.

The experimentally derived X-ray structure revealed unexpected deviations from the design model. While the Rosetta structure prediction and design methods themselves are only approximating, the observed differences might arise from other factors as well. Nonetheless, we were able to insert a new element into the TIM barrel without disrupting its structural integrity or stability. The newly inserted 3^10^ helix appears to fit even more snugly on top of the barrel, and the tight packing of the Met94 side chain with the hydrophobic core of the barrel might have caused the observed structural adjustments that even cause the subsequent barrel helix to take an alternate conformation illustrating the plasticity of this *de novo* TIM barrel.

This work shows how idealized protein designs can be diversified and provides a first building block for further designs towards sTIM11-based enzymes. Due to the four-fold symmetry of sTIM11, this extension can even be transferred to any of the related βα-loops. It therefore provides a flexible extension that might as well be combined with other structural modifications helping to form binding or catalytic sites.

## Methods

### Computational methods

PSIPRED (version 4.0) was used to predict the secondary structure of randomly generated sequences as candidates for the barrel extension. The Rosetta molecular modeling suite (weekly release, first quarter 2016) was used for *ab initio* folding of the extension containing barrel fragment by using the *ab initio* relax method. The Rosetta relax method was used to smooth structural deviations from manual intervention, when transferring the folded helix extension to the entire barrel, which was performed using PyMOL (version 1.8). Finally, the Rosetta enzyme design method was used to adjust the helix residues to adjacent barrel residues.

### Cloning methods

In order to insert the gene fragment encoding the extension, we generated two overlapping fragments in a first PCR reaction. Both samples were loaded on a 1 % agarose gel and after electrophoresis, appropriate bands were excised and purified with the PCR clean-up kit (Qiagen). These two purified DNA fragments served as templates for the second PCR reaction, where both fragments were combined at their overlapping region. The PCR sample was loaded on a 1 % agarose gel and after electrophoresis, the DNA band with the correct size was excised and purified as before.

Purified DNA fragment and vector pET21b(+) were then double-digested using *NdeI* and *XhoI* (Fermentas) at 37 °C for 2 hours. After digestion, the cut vector was purified using the PCR clean-up kit (Qiagen), while the DNA fragment was purified and in parallel concentrated using the DNA Clean & Concentrator Kit (Zymo Research). Vector and fragment were ligated in an overnight reaction at 4 °C using T4 DNA Ligase (New England Biolabs). After transformation, successful clones were verified by sequencing.

### Protein expression and purification

Transformants of *E. coli* BL21 were grown in LB media at 37 °C in a shaker until the culture reached an OD_600_ of 0.6. Protein expression was induced by addition of IPTG (1 mM). The culture was incubated at 37°C for another 4 hours and then harvested by centrifugation. After washing and resuspension in buffer A (50 mM potassium phosphate buffer pH 8.0, 150 mM NaCl, 20 mM Imidazole), cells were lysed by pulsed sonication. Soluble and insoluble components were separated by centrifugation (20000 *g*).

### Biochemical characterization

The biochemical characterization was done in potassium phosphate buffer (50 mM pH 8.0, 150 mM NaCl) using a sample concentration of 0.2 mg mL^−1^. For circular dichroism (CD), the sample was placed in a 1 mm cuvette and the spectrum was recorded from 240-195 nm on a Spectropolarimeter (Jasco J-810). Measured values were normalized to the molar ellipticity per amino acid.

The melting temperature was determined using the same sample. CD signal was tracked at 222 nm, while the instrument heated up or cooled down respectively from 30 °C to 95 °C with a rate of 1 K min^−1^. Measured values were transformed to display the fraction of folded and unfolded protein.

Analytical size-exclusion chromatography was done with a Superdex 75 10/300 GL (GE Healthcare Life Sciences) on an ÄKTApurifier system (GE Healthcare Life Sciences). The sample (0.5 mL) was loaded on the buffer equilibrated column and the absorption was tracked at 280 nm.

### Structure determination

Crystallization conditions were screened using commercially available sparse-matrix screens (Qiagen) and a protein concentration of 4.7 mg mL^−1^. Sitting drops (1 μL) in a ratio of 1:1 (protein:screening solution) were pipetted into a 3-well Intelli-Plate (Art Robbins Instruments) using a Honeybee 961 (Genomic solutions). Plates were sealed with clear tape and incubated at 20 °C. Crystals were obtained in a condition containing 0.2 M Lithium sulfate, 0.1 M Tris pH 8.5, and 30% PEG 4000. Crystals were mounted and flash-cooled in liquid nitrogen. Diffraction images were collected at 100 K on a Pilatus 2M-F at the beamline X06DA (PX III, Swiss Light Source, PSI). 1400 images were collected using an oscillation of 0.1° per image.

Collected data were processed using XDSAPP 2.0 (Sparta et al., 2016). Phases were solved by molecular replacement using PhaserMR with a quarter of sTIM11 (PDB: 5BVL) as search model (McCoy et al., 2007). Refinements were done using Phenix and manual model building was performed with Coot (Adams et al., 2010; Emsley et al., 2010).

## Data availability

Coordinate and structure files have been deposited to the Protein Data Bank (PDB) with accession code 7A8S.

## Acknowledgment

We acknowledge financial support and allocation of beamtime by HZB and thank the beamline staff at BESSY for assistance. We further thank Kaspar Feldmeier for detailed explanations about the design and structure of sTIM11, and Horst Lechner, André C. Stiel and Sergio Romero-Romero for helpful comments on the manuscript.

## Author contribution

Conceptualization, G.W. and B.H.; Investigation, G.W. and S.S.; Data Curation, G.W. and S.S.; Writing – Original Draft, G.W., S.S. and B.H.; Writing – Review & Editing, G.W., S.S and B.H.; Funding Acquisition, B.H.; Project Administration, B.H.

