## Supplementary Information for "Extension of a *de novo* TIM barrel with a rationally designed secondary structure element"

**Figure S1:** Biochemical characterization of all four variants in comparison
**Table S1:** Amino acid sequences of the design target sTIM11 and the computationally derived variants

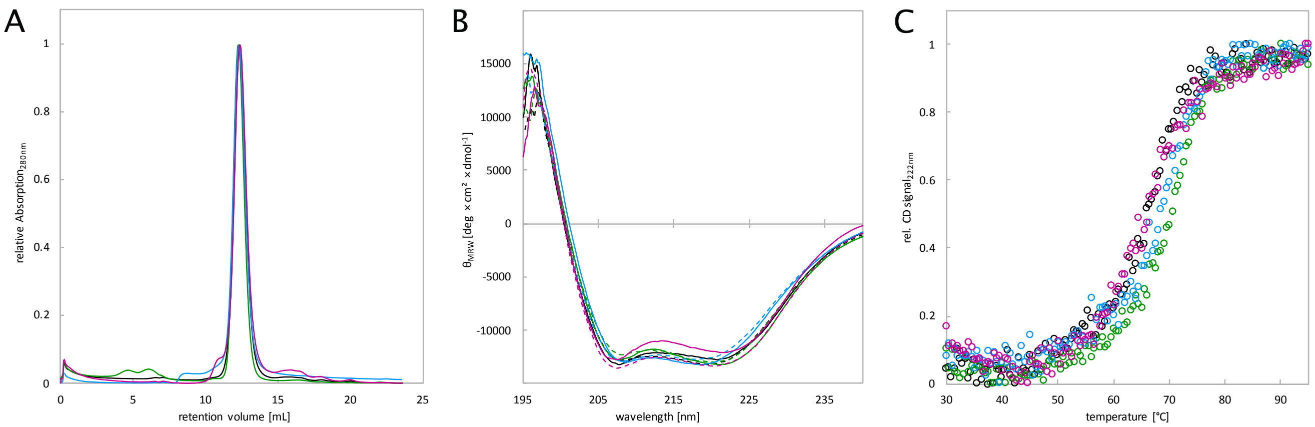


**Supplementary Figure 1: Biochemical characterization of all four variants in comparison.** (A) Analytical size-exclusion chromatography. (B) Far-UV CD before (solid line) and after thermal denaturation (dashed line). (C) Thermal melting measurements. In all three panels sTIM11_helix1 is shown in green, sTIM11_helix2 in blue, sTIM11_helix3 in black, and sTIM11_helix4 in magenta.

**Supplementary Table 1: Amino acid sequences of the design target sTIM11 and the computationally derived variants.** sTIM11 was modified by removing the symmetry breaking cysteines in the sequence (red) that did not form the expected stabilizing disulfide bond in the original work (Huang et al., 2016). Extensions are highlighted in yellow. The third variant yielded an X-ray structure and is presented in this work.

| | |
|---|---|
| sTIM11 (Cysteines removed) | MDKDEAWKQV EQLRREGATQ IAYRSDDWRD LKEAWKKGAD ILIVDATDKD EAWKQVEQLR REGATQIAYR SDDWRDLKEA WKKGADILIV DATDKDEAWK QVEQLRREGA TQIAYRSDDW RDLKEAWKKG ADILIVDATD KDEAWKQVEQ LRREGATQIA YRSDDWRDLK EAWKKGADIL IVDATGLEHH HHHH |
| sTIM11 _helix 1 | MDKDEAWKQV EQLRREGATQ IAYRSDDWRD LKEAWKKGAD ILIVDATDKD EAWKQVEQLR REGATQIAYR SDDWRDLKEA WKKGADILIV DQAEMMQNGM SKDEAWKQVE QLRREGATQI AYRSDDWRDL KEAWKKGADI LIVDATDKDE AWKQVEQLRR EGATQIAYRS DDWRDLKEAW KKGADILIVD ATGLEHHHHH H |
| sTIM11 _helix 2 | MDKDEAWKQV EQLRREGATQ IAYRSDDWRD LKEAWKKGAD ILIVDATDKD EAWKQVEQLR REGATQIAYR SDDWRDLKEA WKKGADILIV DEAQMRQNNM PKDEAWKQVE QLRREGATQI AYRSDDWRDL KEAWKKGADI LIVDATDKDE AWKQVEQLRR EGATQIAYRS DDWRDLKEAW KKGADILIVD ATGLEHHHHH H |
| sTIM11 _helix 3 | MDKDEAWKQV EQLRREGATQ IAYRSDDWRD LKEAWKKGAD ILIVDATDKD EAWKQVEQLR REGATQIAYR SDDWRDLKEA WKKGADILIV SEEMARHAPK DEAWKQVEQL RREGATQIAY RSDDWRDLKE AWKKGADILI VDATDKDEAW KQVEQLRREG ATQIAYRSDD WRDLKEAWKK GADILIVDAT GLEHHHHHH |
| sTIM11 _helix 4 | MDKDEAWKQV EQLRREGATQ IAYRSDDWRD LKEAWKKGAD ILIVDATDKD EAWKQVEQLR REGATQIAYR SDDWRDLKEA WKKGADILIV GDAKQCRQKG LKDEAWKQVE QLRREGATQI AYRSDDWRDL KEAWKKGADI LIVDATDKDE AWKQVEQLRR EGATQIAYRS DDWRDLKEAW KKGADILIVD ATGLEHHHHH H |
